## Supplementary Figure S.1 to S.5 for "Probing the Link Between Vision and Language in Material Perception Using Psychophysics and Unsupervised Learning"

**Psychophysical Stimuli.**
We sampled 72 images from the Space of Morphable Material Appearance. The images were generated from our StyleGAN models. Top three rows are the images of 'Original' materials: Soap, Toy, and Rock. They were synthesized from the trained StyleGAN generators: $G_{soap}$, $G_{toy}$, and $G_{rock}$, respectively. The bottom three rows are the images of 'Morphed' materials: Soap-to-rock, Rock-to-toy, and Soap-to-toy. They were generated from the morphed StyleGAN generator: $G_{soap\text{-}to\text{-}rock}$, $G_{rock\text{-}to\text{-}toy}$, and $G_{soap\text{-}to\text{-}toy}$, respectively.

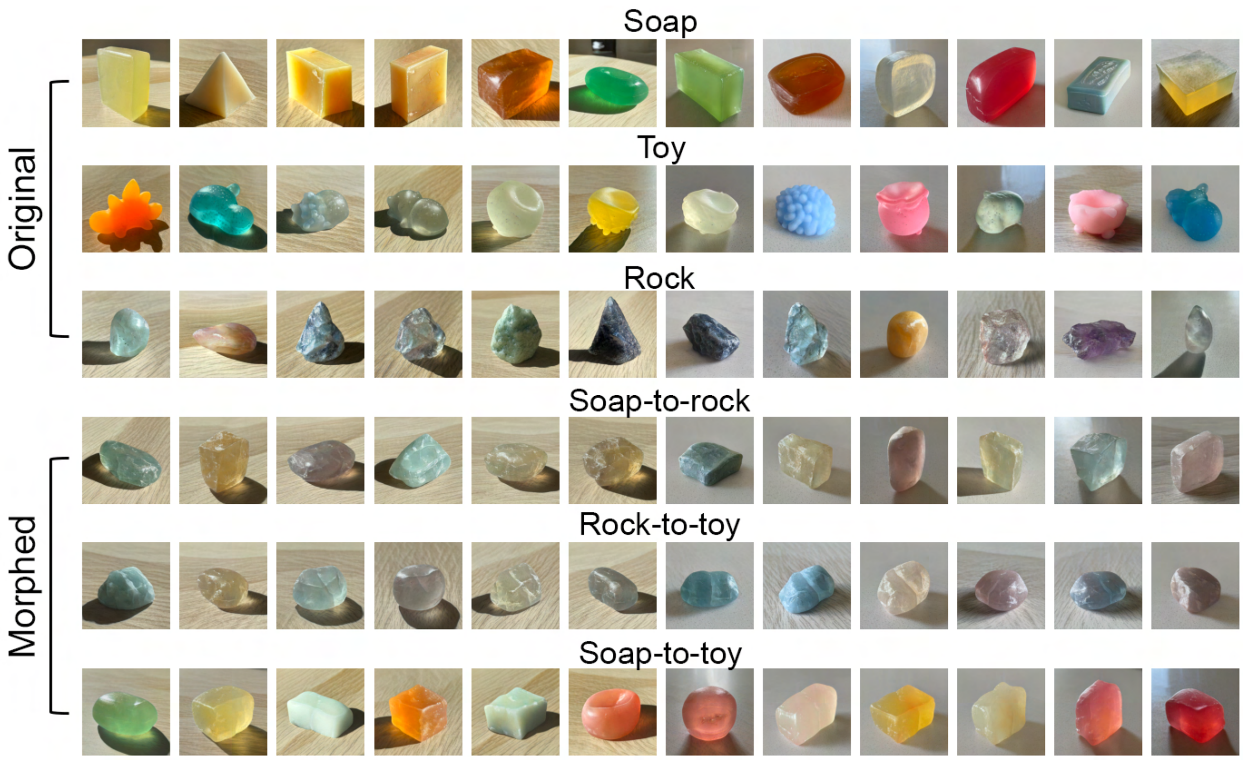


**Figure. S.1.** Seventy-two generated images used as stimuli for both Multiple Arrangement and Verbal Description experiments.

**Perceptual Representational Dissimilarity Matrices (RDMs) from psychophysical experiments.**
Here, we present the individual participant's (N=16) Vision RDMs (from the Multiple Arrangement experiment) and Text RDMs (from the Verbal Description experiment).

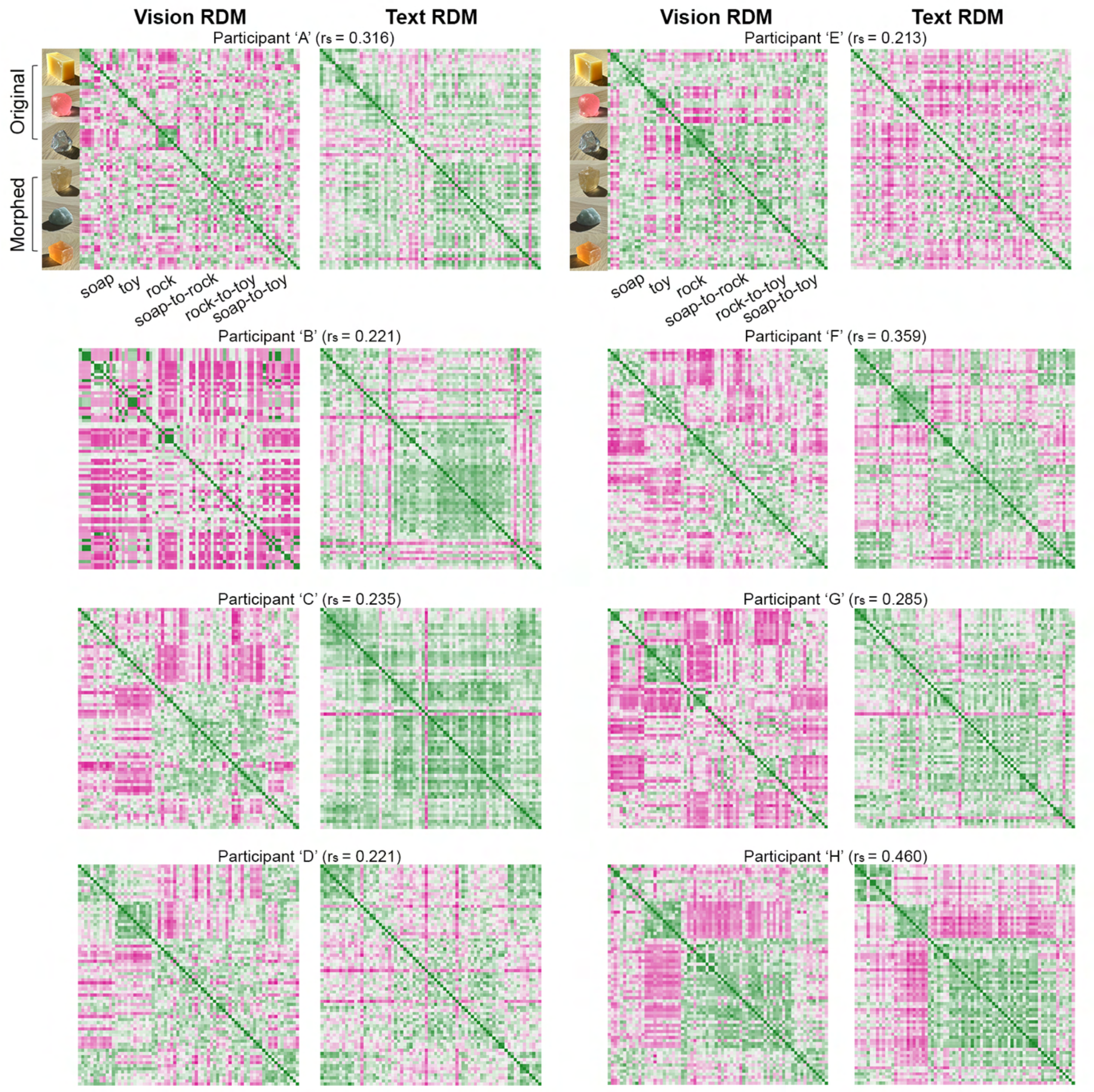

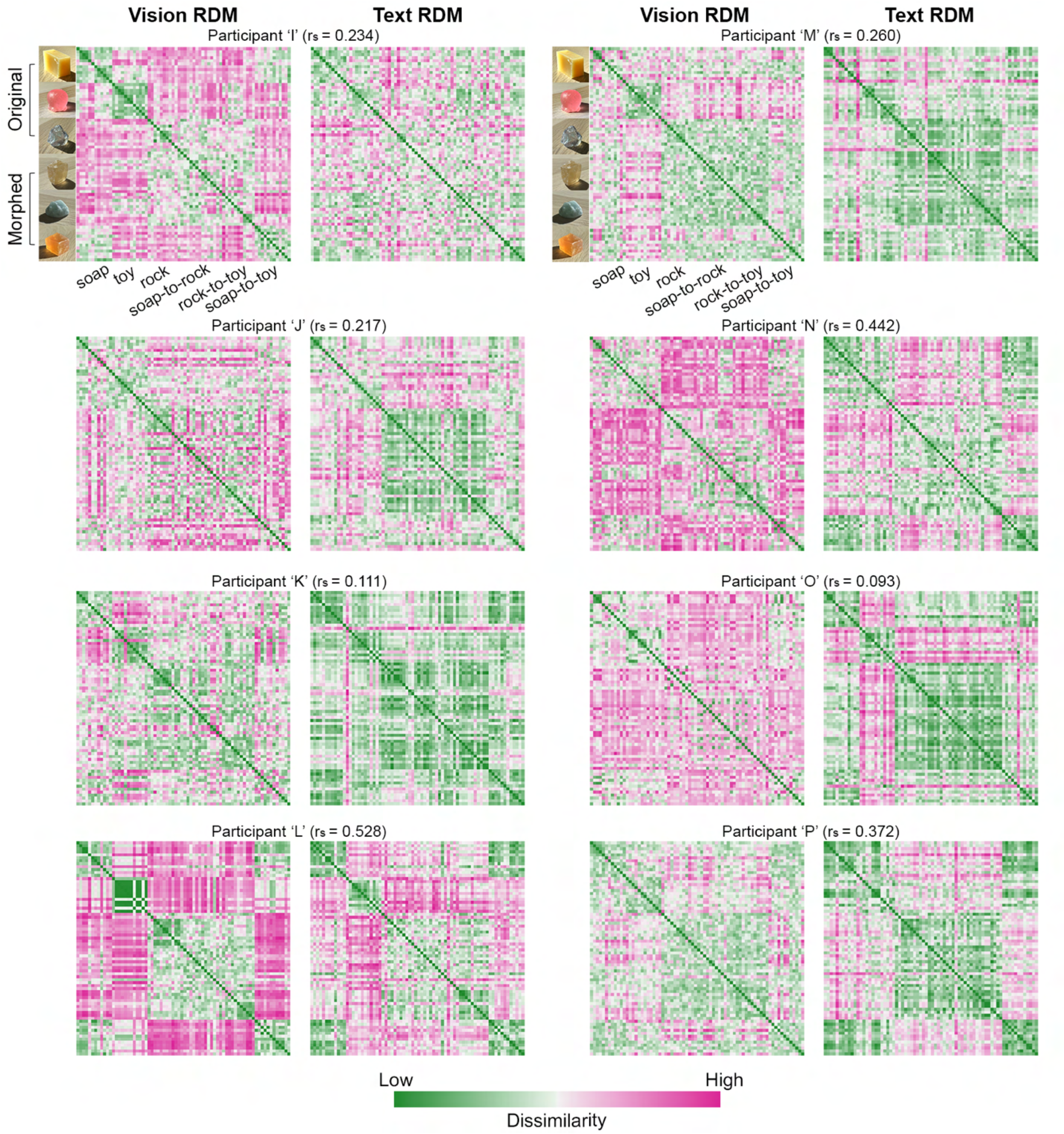


**Figure S.2.** Individual participant's (N=16) RDMs of visual material similarity judgment via Multiple Arrangement (Vision RDMs) and Verbal Description (Text RDMs). The Text RDMs are based on the CLIP's text embedding results, as illustrated in the main paper Figure 4A. The Spearman's correlation ($r_s$) between the participant's own Vision and Text RDMs is marked on top of each pair of RDMs.

**Gromov-Wasserstein Optimal Transport (GWOT).**

We applied the unsupervised alignment method GWOT to compare two similarity structures between the stimulus-level alignment between the Vision and Text RDMs. Here, we used the OpenAI's Embedding V3 to construct the individual participant's Text RDM and compute the group average Text RDM.

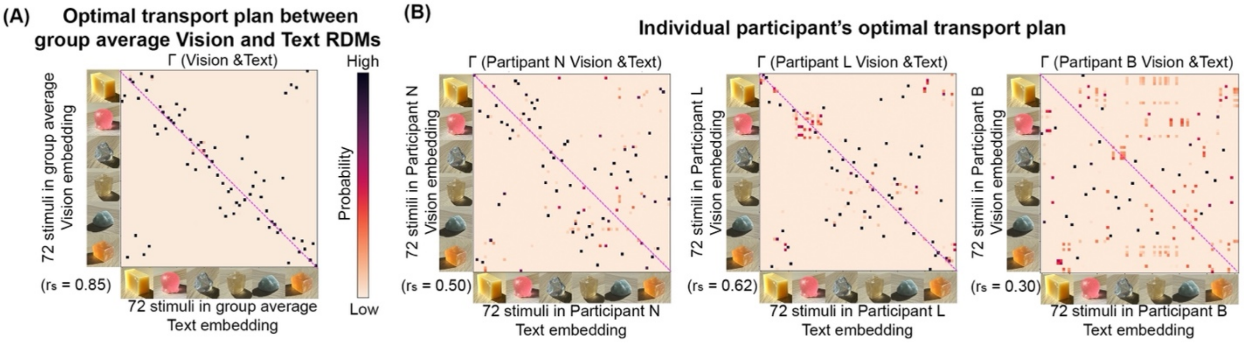


**Figure S.3.** Optimal transportation plan between Vision and Text RDM. The Text RDMs are based on OpenAI's Embedding V3. (A) Optimal transportation plan matrix (Γ) between group average Vision and Text RDMs. (B) Optimal transportation plan matrix of individual participant's Vision and Text RDMs. The Spearman's correlation ($r_s$) between the Vision and Text RDMs is noted in the bottom left corner of the Γ matrix.

**Image representations from the pre-trained models.**

We tested various visual-semantic models, self-supervised vision models, and perceptual similarity metrics. For the visual-semantic models, we examined the embeddings resulting from various versions (using either ResNet or Vision Transformer (ViT) as the backbone) of their image encoder: CLIP-ResNet50, CLIP-ViT-B/16, CLIP-ViT-B/32, CLIP-ViT-L/14, OpenCLIP-ViT-B/32, OpenCLIP-ViT-L/14, and OpenCLIP-ViT-H/14. For the self-supervised vision models, we tested the family of DINO: DINO (e.g., DINO-S8) and DINOV2 (DINOV2-small [1]). We evaluated the image similarity based on two established metrics: the lower-level patch-based metric LPIPS, and the more semantically aware metric DreamSim [2].

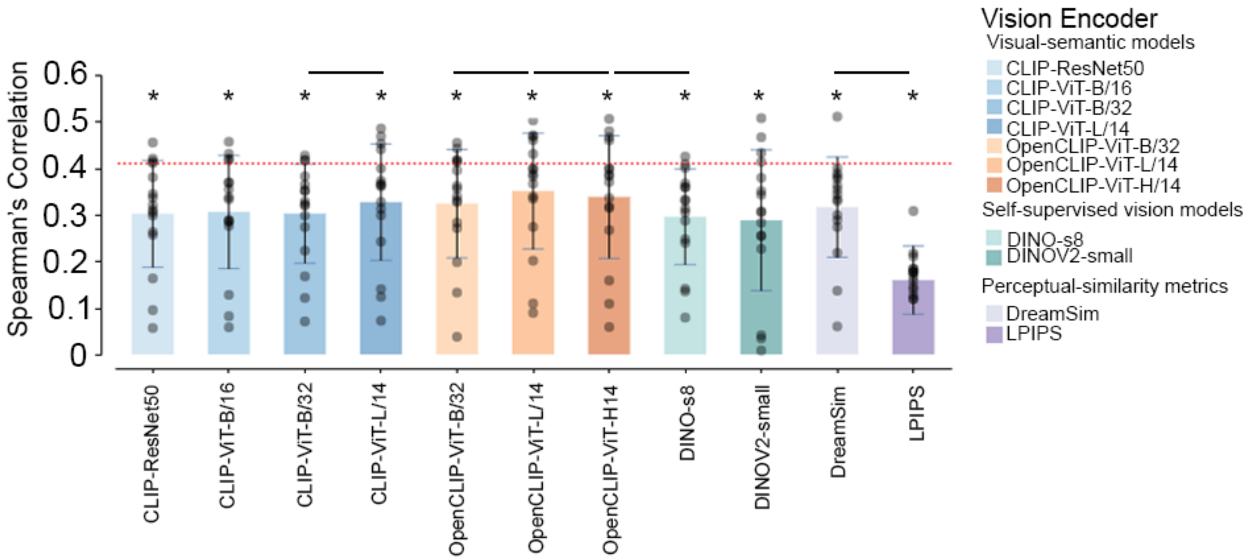


**Figure S.4.** Spearman's correlation between an individual's Vision RDM and Image-feature RDM from each tested vision encoder. The bars represent the average correlations across participants. The block dots represent the individual participants. The red dotted line indicates the lower bounds of the noise ceiling of human visual judgment results. On top of each bar, * indicates $p < 0.005$ for model-specific one-sided signed-rank tests against zero. The horizontal black bar indicates $p<0.05$ for two-sided pairwise signed-rank tests between two nearby vision encoder models shown in the plot.

**Space of Morphable Material Appearance.** Given two images of a pair of source and target materials, we can produce a morphing sequence that smoothly the material appearance in the image space. With a pair of source and target generators, $G_{source}$ and $G_{target}$, we apply linear interpolation to the models' weights at all convolution layers, while also interpolating between the latent codes drawn from the corresponding latent spaces $W_{source}$ and $W_{target}$ (see Method in main paper). We can synthesize the corresponding morphed material at any interpolation step, $\lambda$.

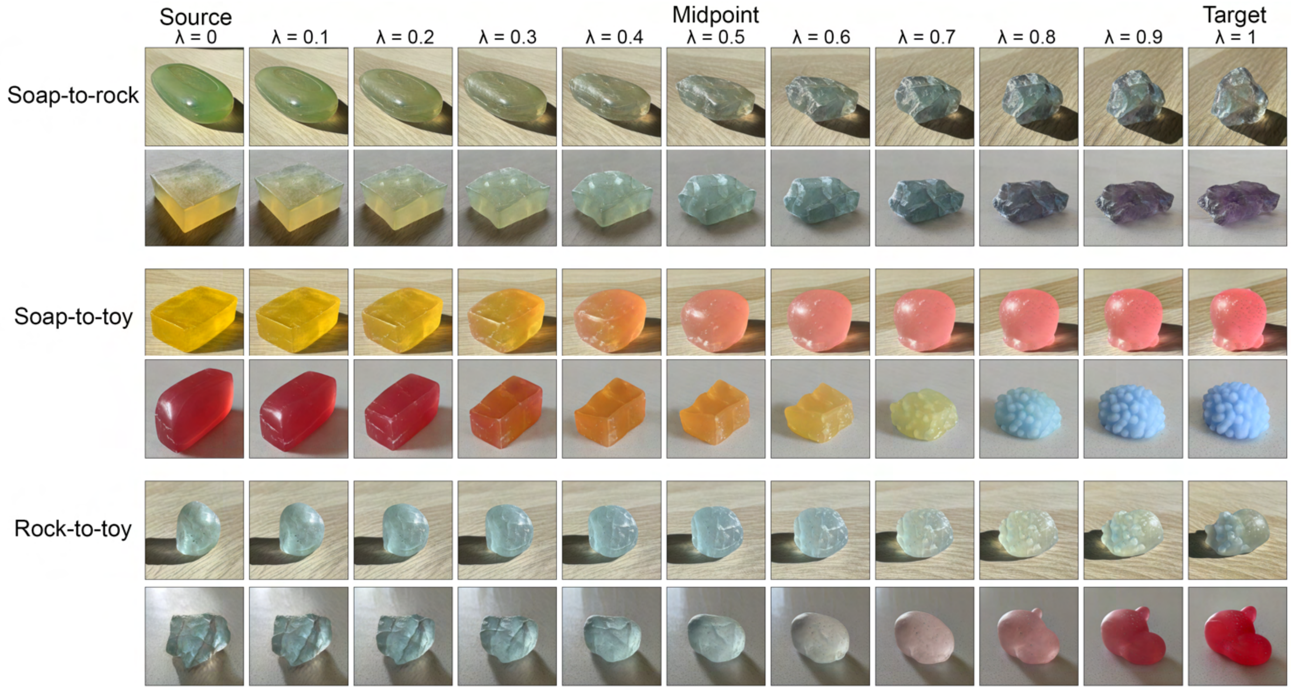


**Figure S.5.** Cross-material morphing. The source material transforms into the target material with a nine-step interpolation.
